# The glucocorticoid receptor in the nucleus accumbens plays a crucial role in social rank attainment in rodents

**DOI:** 10.1101/668897

**Authors:** Aurélie Papilloud, Meltem Weger, Alexandre Bacq, Ioannis Zalachoras, Fiona Hollis, Thomas Larrieu, Dorian Battivelli, Jocelyn Grosse, Olivia Zanoletti, Sébastien Parnaudeau, François Tronche, Carmen Sandi

**Author notes:** Correspondence: Professor C Sandi, Laboratory of Behavioral Genetics, Brain Mind Institute, Ecole Polytechnique Fédérale de Lausanne, Station 19, CH-1015 Lausanne, Switzerland. Equal senior contribution (François Tronche, Carmen Sandi).

## Abstract

Social hierarchy in social species is usually established through competitive encounters with conspecifics. It determines the access to limited resources and, thus, leads to reduced fights among individuals within a group. Despite the known importance of social rank for health and well-being, the knowledge about the processes underlying rank attainment remains limited. Previous studies have highlighted the nucleus accumbens (NAc) as a key brain region in the attainment of social hierarchies in rodents. In addition, glucocorticoids and the glucocorticoid receptor (GR) have been implicated in the establishment of social hierarchies and social aversion. However, whether GR in the NAc is involved in social dominance is not yet known. To address this question, we first established that expression levels of GR in the NAc of high anxious, submissive-prone rats are lower than that of their low anxious, dominant-prone counterparts. Furthermore, virally-induced downregulation of GR expression in the NAc in rats led to an improvement of social dominance rank. We found a similar result in a cell-specific mouse model lacking GR in dopaminoceptive neurons (i.e., neurons containing dopamine receptors). Indeed, when cohabitating in dyads of mixed genotypes, mice deficient for GR in dopaminoceptive neurons had a higher probability to become dominant than wild-type mice. Overall, our results highlight GR in the NAc and in dopaminoceptive neurons as an important regulator of social rank attainment.

## Introduction

Social rank influences behavior and physiology in both humans and animals, and has been linked with the development of psychopathologies (1,2). Most social species are organized in social hierarchies. The organizing principle of social hierarchies is to provide dominant individuals with priority access to resources, such as new territory, food, water and mating partners (3,4).

In laboratory rodents, social hierarchy is often established through repeated agonistic and antagonistic or aggressive interactions. It usually develops within a few days and remains stable over long periods of time (5). The reward- and motivation-related mesolimbic dopamine system has been identified to be important for the establishment of social dominance [reviewed in (6)]. According to previous observations in rodent models for trait anxiety, the nucleus accumbens (NAc) is actively engaged by aggressive encounters. Furthermore, metabolic activity in this region has been directly linked with social hierarchy in rat (7,8). Specifically, dopaminoceptive D1 receptor-containing neurons in the NAc are activated by social competition, particularly in dominant-prone individuals (8,9), while blockade of D1 receptors diminished social dominance (9). Altogether, these observations indicate an important role for NAc dopaminoceptive neurons in the establishment of social rank.

Several lines of evidence indicate that stress response and more particularly glucocorticoid hormone may also play a key role in the establishment of hierarchy. We have previously shown that acute stress increases the propensity to become subordinate in a pair of unfamiliar rats matched for age, body weight and anxiety levels (10). Glucocorticoids levels rise in the bloodstream upon social defeat (11), following social encounters (12,13) and have been recently shown to participate to the establishment and maintenance of social hierarchies (14,15). Importantly, selective inactivation of the gene encoding the glucocorticoid receptor (GR) in dopamine-innervated brain regions (including the striatum, the NAc and the deep cortical layers) was shown to prevent social aversion in animals undergoing social defeat (11). Altogether these data indicate that glucocorticoids through GR might play a key role in shaping behavioral trajectories leading to social rank attainment. However, to date it is not known whether GR in the NAc is important for the modulation of social hierarchies.

In this study, we investigated the potential role of GR in the NAc and dopaminoceptive neurons in rank attainment. To this end, we first analyzed expression levels of GR and mineralocorticoid receptor (MR) in the NAc in high and low anxious rats that are, respectively, prone to become submissive or dominant, when submitted to dyadic encounters in competition for a new territory. Next, we explored the causal involvement of GR in rank attainment through adeno-associated virus (AAV)-induced GR downregulation in the NAc of rats. Finally, in order to address GR involvement in a cell type specific manner, we investigated rank attainment in dyads of cohabitating mice involving one wild-type mouse and a mouse lacking GR in dopaminoceptive neurons.

## Materials and Methods

### Animals

Adult male Wistar rats (Charles River) weighing 250-275 g at the start of experiments were used. For the initial experiments that examined social hierarchy between high and low anxious animals, rats were singly housed upon arrival to the vivarium to avoid any influence from social experiences. For all other experiments, after arrival, animals were housed 2 per cage and allowed to acclimate to the vivarium for one week. All animals were subsequently handled for 2 minutes for a minimum of 3 days. They were weighed upon arrival as well as weekly to ensure good health.

Mouse experiments were conducted in weight-matched male GR^D1Cre^ mice and their control littermates bred in a C57Bl/6 background. The generation of GR^D1Cre^ mice has been described previously (16). Briefly, Nr3c1 (*GR*) gene inactivation was selectively targeted in dopaminoceptive neurons (Nr3c1^loxP/loxP^;(Tg)D1aCre (17) hereafter designed GR^D1Cre^). Experimental animals were obtained by mating Nr3c1^loxP/loxP^ females with Nr3c1^loxP/loxP^;Tg:D1aCre males.

Rats and mice were maintained under standard housing conditions on a 12h light-dark cycle (lights on at 7:00 AM). Food and water were available ad libitum. Experiments on rats were performed with the approval of the Cantonal Veterinary Authorities (Vaud, Switzerland) and carried out in accordance with the European Communities Council Directives of 24 November 1986 (86/609EEC). Experiments on mice were performed in accordance with French Ministère de l’Agriculture et de la Forêt (87-848) and the European Directive 2010/63/UE and the recommendation 2007/526/EC for care of laboratory animals.

### Experimental design

One week after their arrival, rats were tested in an elevated plus maze to assess basal anxiety and were classified as high (HA) or low anxious (LA). HA and LA animals were then paired according to their weight and tested for social hierarchy. A separate cohort of HA and LA animals were sacrificed without hierarchy testing and their brains were extracted for gene expression analysis.

For the viral downregulation of GR, animals were assigned to experimental groups with averaged anxiety levels being similar between groups. Rats then underwent surgery for delivery of a viral construct expression shRNA rargeting GR (GR-KD) or a viral construct expressing a scrambled control (SCR) in the NAc (see also Supplementary Materials and Methods). After 6 weeks of recovery, animals’ anxiety levels and locomotion were tested in an open field. SCR and GR-KD rats were then matched by weight and paired in a new cage, avoiding previous cage mates to be placed together. After two weeks of cohabitation, the social confrontation tube test and water competition test were performed. Seventy-two hours later, they were sacrificed and their brains were extracted and used to assess virus localization and efficiency.

Finally, for the experiment involving GR mutant mice, 12 weeks old control and GR^D1Cre^ mice were matched by weight and paired in a new cage. As for rat experiments, pairing of previous cage mates was avoided. After one week of cohabitation, social dominance was assessed in the dyad using the social confrontation tube test and the warm spot test. Seventy-two hours later, mice were sacrificed for basal corticosterone analysis.

### Elevated plus maze test

Rats were tested for anxiety-like behavior in the elevated plus maze (18). For more information, see Supplementary Materials and Methods.

### Open field and novel object reactivity tests

The open field and novel object reactivity tests were performed to assess the rats’ emotional and exploratory/locomotive behavior, as well as their reactivity upon novelty exposure. The tests were performed as previously described (19). For additional details, see Supplementary Materials and Methods.

### Social dominance tests

Dominance was assessed using the social hierarchy test (8), the social confrontation tube test, the water competition test and the warm spot test (20–22). For further details, see Supplementary Materials and Methods.

### Viral downregulation of GR

Two weeks after arrival, rats were subjected to surgery for the viral downregulation of GR in the nucleus accumbens, wherein an adeno-associated AAV1/2 vector containing an U6-pm-GR3 shRNA-terminator-CAG-EGFP-WPRE-BGH-polyA-expression cassette was used. Control animals were injected with a scrambled shRNA construct (AAV1/2-U6-SCR shRNA-CAG-EGFP-WPRE-BGH-polyA). All viral constructs used were designed and produced by GeneDetect, New Zealand. For further details, see Supplementary Materials and Methods.

### Corticosterone analysis

Trunk blood was collected during sacrifice for basal corticosterone measurement in both rats and mice. Animals were sacrificed in the morning. Free corticosterone measurements were obtained from blood plasma samples via centrifugation and subsequent measurement using an enzymatic immunoassay kit that was performed according to manufacturer’s instructions (Enzo Life Sciences, Switzerland). Levels were calculated using a standard curve method.

### Statistical analysis

Data were analyzed with Student *t*-test or Mann-Whitney test, as appropriate, using the statistical package GraphPad Prism 5 (GraphPad software Inc., USA). If Levene’s test for equality of variances was significant, equal variance was not assumed and the altered degree of freedom was rounded to the nearest whole number. Dominance in the tube test was analyzed with Fisher’s exact test. The Pearson’s chi-squared test was used to analyze the independence of the tube test and the warm spot test. All bars and error bars represent the mean ± SEM. Significance was set at *p* < 0.05, while the *p*-values were considered tending toward significance when 0.05 ≤ *p* ≤ 0.1. Graphs were created using GraphPad Prism 5.

## Results

### Submissive-prone rats show reduced glucocorticoid receptor gene expression in the nucleus accumbens

As reported previously (8,9), high anxious rats had a higher probability to lose a competition for a new territory when confronted with low anxious counterparts. To understand if the outcome of the social competition could be considered as a phenotypic trait, we analyzed the hierarchical behavior across numerous cohorts of high and low anxious animals. Qualitative analyses of offensive behaviors indicated that high anxious rats exhibited reduced offensive behavior both for the compound index (Figure 1b; *U* = 1861, *p* < 0.0001) and individually, for each of the respective offensive behaviors (Figure 1a, b; offensive upright: *U* = 2741, *p* < 0.0001; keeping down: *U* = 2308, *p* < 0.0001; lateral threat: *U* = 2912, *p* < 0.0001). Therefore, we consider high anxious animals as submissive-prone (subP) and low anxious rats as dominant-prone (domP) animals.

**Figure 1.**
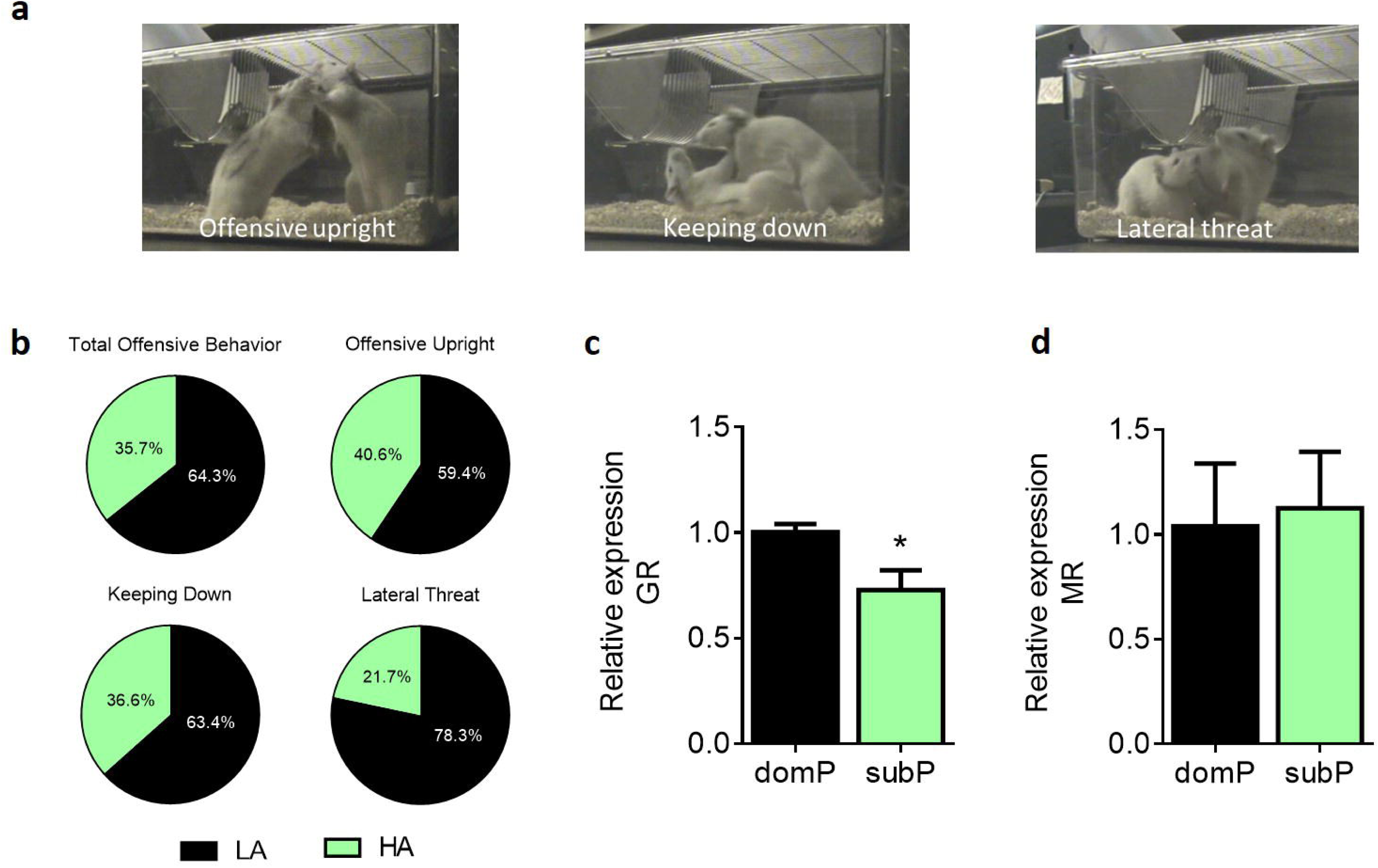
Characterization of dominant and submissive-prone rats in behavior and gene expression. Example of rats illustrating the three dominant postures considered in the social hierarchy test: offensive upright, keeping down and lateral threat **(a).** High anxious (HA) rats exhibited lower percentage of total offensive behavior, offensive upright, keeping down and lateral threat in a social hierarchy test than low anxious (LA) animals **(b).** GR expression analysis in the nucleus accumbens (NAc) revealed lower expression levels in submissive-prone (subP) rats in comparison to dominant-prone (domP) animals **(c).** The mRNA levels of MR were similar between groups. N: LA/domP rats = 97 (b) or 4-5 (c, d) and LA/subP rats = 97 (b) or 4-5 (c-d). *p < 0.05, vs domP. Results are expressed as mean ± SEM.

We next assessed the expression levels of GR and MR genes in the NAc in an independent group of rats, which were only tested for anxiety levels. We found significantly lower GR gene expression in the NAc of subP rats compared to domP conspecifics (Figure 1c; *t*_7_ = 2.93, *p* < 0.05). In contrast, MR expression was similar for both groups (Figure 1d; *t*_7_ = 0.45, *p* = 0.67). Interestingly, basal corticosterone levels were higher in subP rats than in domP animals (Figure S1).

### Downregulation of the glucocorticoid receptor in the nucleus accumbens modifies spontaneous behaviors in novel environments

In order to investigate whether GR in the NAc plays a role in social competition, we injected an adeno-associated virus downregulating GR (GR-KD) or, in the control group, a virus expressing a scrambled construct (SCR) into the NAc (Figure S2a). Knockdown efficiency in the NAc was confirmed through qRT-PCR quantification of GR mRNA levels (Figure S2b) and by immunostaining (Figure S2c). Six weeks after the injection, anxiety-like and exploratory behaviors were assessed in the open field (OF) and novel object reactivity (NOR) tests. In the OF test, GR-KD rats spent a higher percentage of time in the center (Figure 2b; *U* = 44.00, *p* < 0.05), and less time in the wall area (Figure 2c; *t*_25_ = 3.10, *p* < 0.01) than SCR rats. The distance traveled during the test was significantly longer in GR-KD animals (Figure 2d; *t*_25_ = 3.66, *p* < 0.01). Similarly, in the NOR test, GR-KD rats spent more time in the center (Figure 2e; *U* = 44.00, *p* < 0.05) and, consequently, less time in the wall area (Figure 2f; *t*_25_ = 2.44, *p* < 0.05) than SCR animals. Moreover, there was a trend for GR-KD rats to move greater distance than SCR-infused controls (Figure 2g; *t*_25_ = 1.83, *p* = 0.08). Finally, GR-KD rats spent more time sniffing the novel object than SCR conspecifics (Figure 2h; *U* = 47.00, *p* < 0.05).

**Figure 2.**
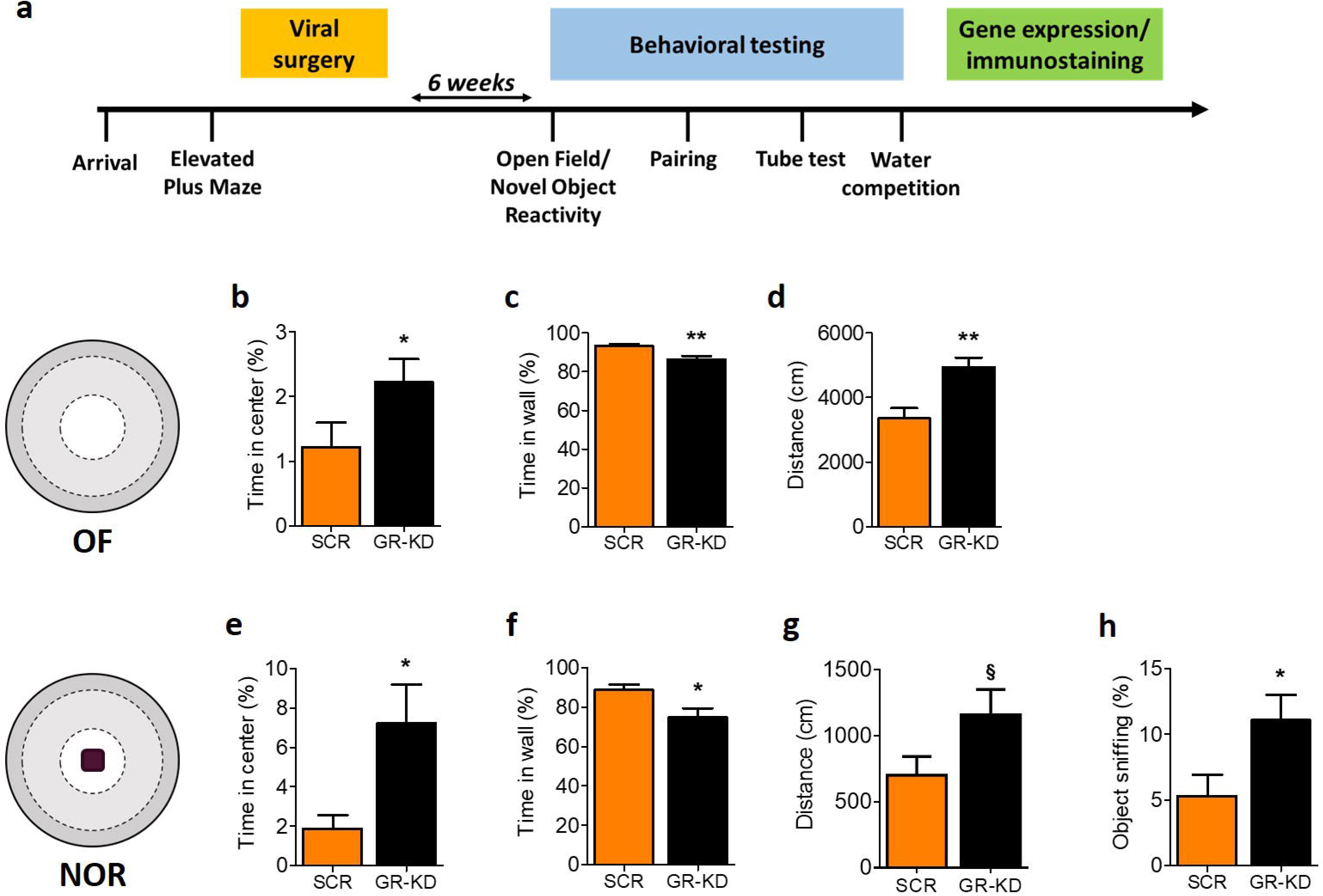
GR knockdown in the nucleus accumbens reduces anxiety-like behavior in the open field and novel object reactivity test. Experimental design **(a)**. GR-KD rats spent more time in the center **(b)** of the open field (OF) and reduced time in the wall zone **(c)** compared to SCR animals. The distance traveled was higher in GR-KD rats **(d).** Similarly, GR-KD rats spent more time in the center when a novel object was presented (NOR) **(e)** and less time in the wall zone **(f)** than SCR animals. The distance moved tended to be higher in GR-KD rats when the object was present **(g).** Furthermore, GR-KD animals spent more time sniffing the object than SCR rats **(h).** *N:* SCR = 12; GR-KD = 14-15. \**p* < 0.05, \*\**p* < 0.01, §*p* < 0.1, vs SCR. Results are expressed as mean ± SEM.

### Downregulation of the glucocorticoid receptor in the nucleus accumbens promotes social dominance

Animals were matched for body weight and placed together to cohabitate in a new cage in dyads, involving one animal from each group (i.e., GR-KD and SCR rats). Two weeks after the beginning of their cohabitation they underwent the social confrontation tube test (Figure 3a). Social rank was considered stable when rats’ performance yielded the same rank for four consecutive days (see Figure 3b for an example from a single cage, Figure 3c for the average of the 12 cages). GR-KD rats had a higher number of winning trials, indicating that they exhibited more dominance than SCR animals (Figure 3d; Fisher’s exact test, *p* < 0.05). In the winning trials, the latency to push the opponent out of the tube was similar for both groups (Figure 3e; *U* = 12.00, *p* = 0.52). The dominance behavior in GR-KD rats was shown in a second test, the water competition test, with GR-KD rats spending more time drinking the water than SCR counterparts (Figure S3). No group difference in basal corticosterone levels was detected (Figure 3f; *U* = 142.00, *p* = 0.73).

**Figure 3.**
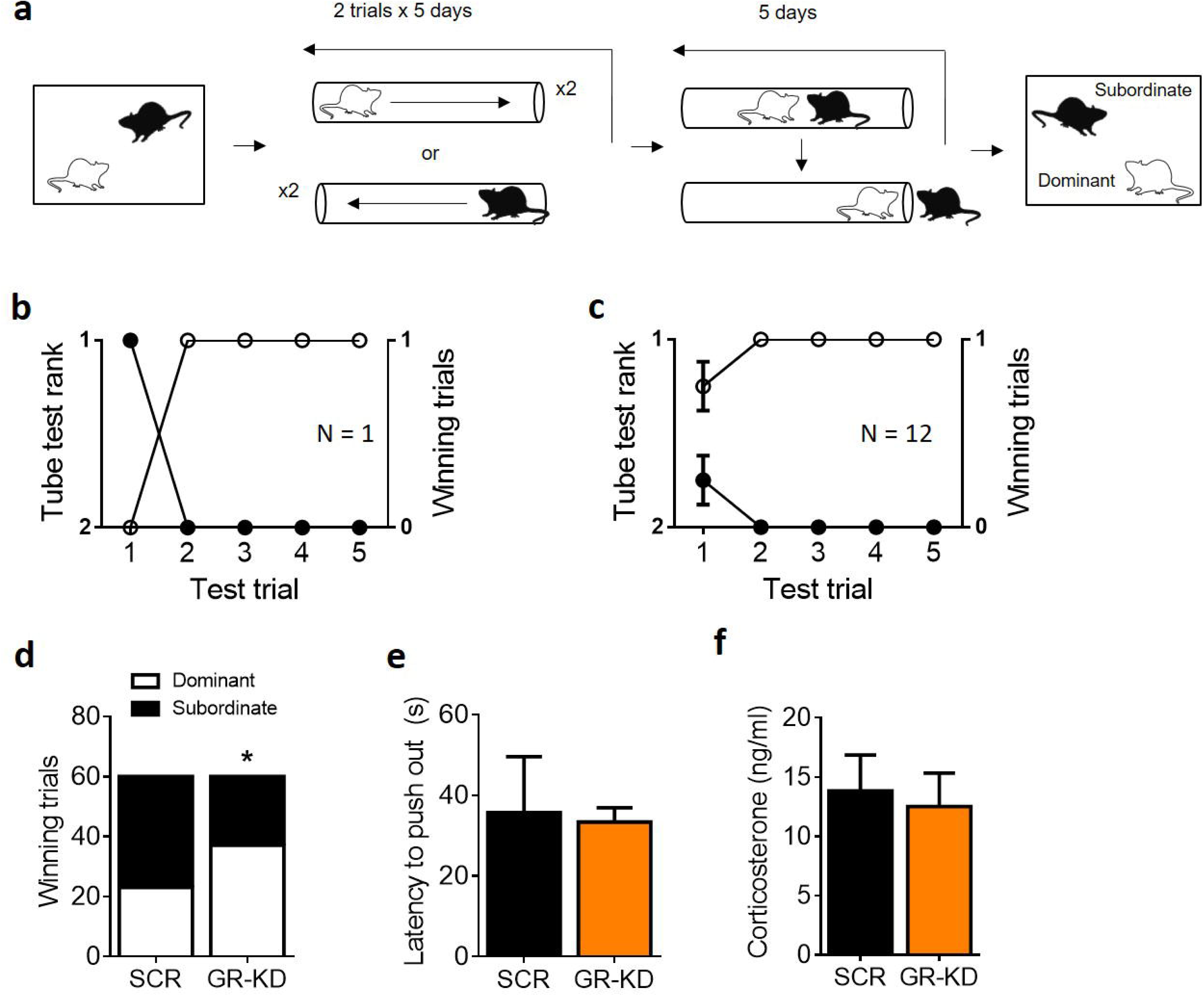
GR knockdown in the NAc enhances social rank attainment in a dyadic hierarchy in rats. Following dyadic cohabitation of a GR-KD and a SCR-infused rats for two weeks, each dyad of rats was tested for dominance using the social confrontation tube test **(a).** Example of one cage representing the rank and winning trials as function of tube test trials **(b).** Summary for twelve cages over the 5-day test trials **(c).** GR-KD rats had a higher number of winning trials than SCR animals **(d).** The latency to push the opponent out of the tube was similar between groups **(e).** Basal corticosterone measurement at sacrifice was not significantly different between groups **(f)** *N:* SCR = 12, GR-KD = 12. \**p* < 0.05, vs SCR. Results are expressed as mean ± SEM.

### Glucocorticoid receptor gene deletion in dopaminoceptive neurons promotes social dominance in mice

Next, we assessed whether the GR knockout specifically in dopaminoceptive neurons affects the emerging social status in mouse dyads. To this end, dyads constituted by one control and one GR^D1Cre^ mice were formed. One week afterwards, the emerged social hierarchy was tested with the social confrontation tube test (Figure 4b). We performed two replication experiments (see figure S4 for the detailed results of each experiment), and data presented here are pooled. We considered social rank as stable when mice performance yielded the same rank for more than seven consecutive trials (see Figure 4c for an example from a single cage, Figure 4d for the average of all tested pairs). Note that the threshold is different from the one in rats, according to a protocol developed in our lab on the model described by Wang and colleagues (20). GR^D1Cre^ mice won the tube contest more times and, therefore, were considered to be more dominant than controls (Figure 4e; Fisher’s exact test, *p* < 0.001). In the winning trials, there were no group differences in the latency to push the opponent out of the tube (Figure 4f; *U* = 953.50, *p* = 0.67). The dominance rank across dyads was confirmed with the warm spot test (21,22) (Figure S5). Basal corticosterone levels were similar between groups (Figure 4g; *U* = 41.50, *p* = 0.22).

**Figure 4.**
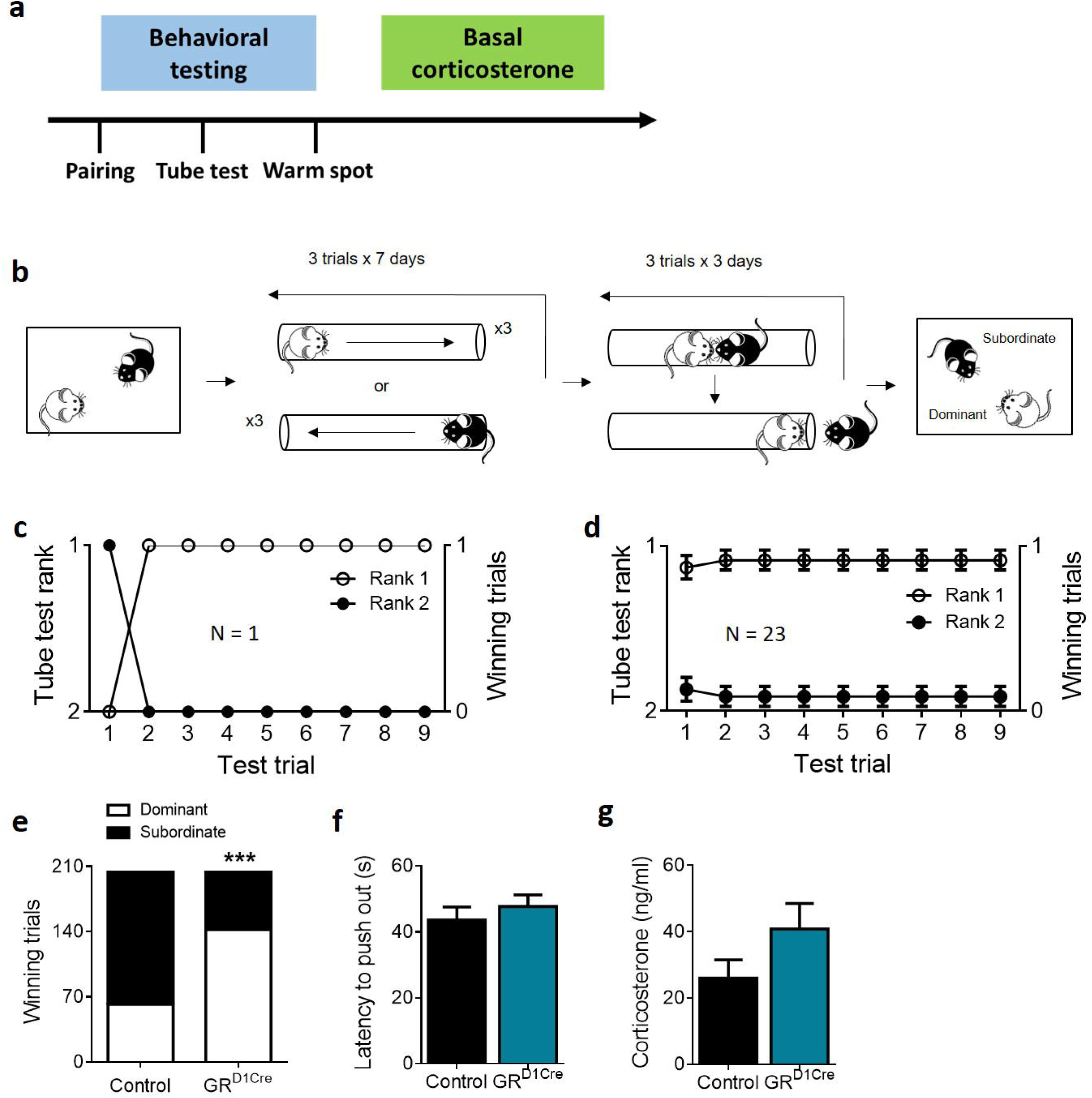
Impact of GR gene inactivation in dopaminoceptive neurons on social rank attainment in a dyadic hierarchy. Experimental design **(a).** Following cohabitation of a dyad consisting of a control and a GR^D1Cre^ mouse for one week, each dyad of mice was tested for dominance using the social confrontation tube test **(b).** Example of one cage representing the rank and winning trials as function of tube test trials **(c).** Summary for 23 cages over the 3-day test trials **(d).** GR^D1Cre^ mice had a higher number of winning trials than control animals **(e).** The latency to push the opponent out of the tube however did not differ **(f).** Finally, basal corticosterone levels measured at sacrifice were similar between groups **(g).** N: Control = 23 (e) or 11 (f, g), GRD1Cre = 23 (e) or 11 (f, g). \*\*\**p* < 0.001, vs control. Results are expressed as mean ± SEM.

## Discussion

Stress and glucocorticoids have been shown to be important modulators for the long-term establishment of social hierarchies-specifically, for social subordination- in rodents (10,14,15). However, these studies have not addressed whether stress or glucocorticoids modulate social dominance via the GR or the MR, nor where in the brain these effects occur. In this study, we detected decreased gene expression of GR, but not MR, in the NAc of high anxious animals that are more prone to lose a social competition and become subordinate. Thus, we investigated the role of GR for social dominance. We observed that knocking down GR in the NAc led to decreased anxiety and made rats more prone to win the contests in the tube test. Furthermore, a similar result was found in another model in which GR is inactivated within dopaminoceptive neurons including the NAc. Indeed, GR^D1Cre^ mice exhibited more dominant behavior than wild-type mice in the tube test. Overall, our results highlight GR in the NAc and in dopaminoceptive neurons as an important regulator for social rank attainment.

Our observations are in line with previous studies reporting a prominent role of the NAc in the development and/or expression of social dominance in rodents (7,8,23–25). Recently, our laboratory has shown that the social competition for a new territory between outbred Wistar rats leads to NAc activation, and that transient pharmacological inhibition of the NAc with the GABA_A_ agonist muscimol results in reduced social competence (8). Similarly, neuroimaging studies in humans found NAc activation under tasks involving manipulations of social status (26,27) or social competition (28). Moreover, the NAc has been described as a key player in the expression of motivated behavior (29,30), that has been associated with social dominance (31,32). However, we did not detect any differences in the time to push the opponent out of the tube between the genotypes during the winning trials in the tube test and the latency to go to drink in the water competition test (data not shown), we can exclude that *a priori* differences in motivation lead to the herein detected changes in social dominance.

Dopamine is well-known for modulating appetitive and aversive motivational processes through binding to D1 or D2 receptors in the NAc (29,30,33). A dyadic competition engages D1-containing neurons in the NAc of rats (8), while antagonizing D1 receptors in this brain region abolishes the animals’ chances to become dominant (9). These data strongly support a key role of dopamine signaling, more specifically within the NAc in the establishment of social hierarchies. Interestingly, glucocorticoids can stimulate dopamine release in the NAc (34–36). Also, the inactivation of GR gene in the whole central nervous system has been shown to decrease dopamine neurons spontaneous firing within the ventral tegmental area (VTA) (16). The GR^D1Cre^ mice we used in our study have been also reported to exhibit a similar decrease of dopamine neurons firing potentially due to an altered feedback from the NAc to the VTA (16,37). In addition, while repeated social defeat is known to induce a long-term increase of dopaminergic activity, that effect is abolished in GR^D1Cre^ mice (11). It is of note that in this mouse model, D1-expressing but also a part of D2-expressing neurons are recombined, precluding us to point at one specific subpopulation’s role in the observed phenotype. However, the combination of our results in this model along with the ones obtained in the NAc GR knockdown experiment in rat strengthen the idea that GR within this brain region might modulate social hierarchy through an effect on the mesolimbic dopamine pathway activity.

Another interesting observation in our study was that the knockdown of GR in the NAc not only promoted dominance, but also decreased anxiety levels in these animals compared to their respective controls. These findings are in line with the literature highlighting anxiety as a crucial factor for social competition and social status in both rodents (8,9,38) and humans (39,40). The fact that we detected lower GR mRNA levels in submissive-prone rats compared to dominant-prone rats, but GR downregulation in the NAc led to reduced anxiety and higher social rank, might appear conflicting at a first glance. However, these results suggest that the impact on social dominance is dependent on glucocorticoid signaling rather than on GR expression itself. For instance, the reduced GR expression in the NAc of high anxious, subP rats might be the natural consequence of higher glucocorticoid input through the GR. In agreement with this possibility, we detected greater levels of corticosterone in high anxious, subP rats than in low anxious, domP animals. Interestingly, corticosterone levels were similar between GR-KD and SCR rats and between control and GR^D1Cre^ mice. One explanation would be that modulating GR expression in the NAc either since birth or at adulthood might impact the hormonal release.

In conclusion, we highlight GR in the NAc and dopaminoceptive neurons as an important player in the establishment of social rank in rodents. These findings add further evidence to the existing literature and provide novel insights into the factors important in the modulation of social dominance.

## Funding and Disclosure

This project has been supported by grants from the Swiss National Science Foundation [31003A-152614 and -176206; NCCR Synapsy (51NF40-158776 and – 185897)], the European Union’s Seventh Framework Program for research, technological development and demonstration under grant agreement no. 603016 (MATRICS), and intramural funding from EPFL to CS. MW was supported by a Marie Curie Intra-European Fellowship for Career Development (H2020-MSCA-IF-2016; EU grant No. 748051). IZ was supported by an EMBO long-term fellowship (ALTF 1537-2015), co-funded by Marie Curie actions (LTFCOFUND2013, GA-2013-609409). This work was also supported by the French Agence Nationale de la Recherche (ANR3053NEUR31438301 and ANR-14-CE35-0029-01), by a Sorbonne University grant (Emergence), the Labex BioPsy, and the Foundation for Medical Research (FRM Equipe grant) to SP and FT. The funding sources had no additional role in study design, in the collection, analysis and interpretation of data, in the writing of the report or in the decision to submit the paper for publication. This paper reflects only the authors’ views and the European Union is not liable for any use that may be made of the information contained therein.

## Acknowledgements

We would like to thank the groups of Prof. Carmen Sandi and Prof. François Tronche for their excellent technical assistance. We would also like to thank the caretakers from the EPFL Centre de PhénoGénomique (CPG) and the animal facilities of the NPS and IBPS.

## References

1. Allan S, Gilbert P. Submissive behaviour and psychopathology - Allan - 2011 - British Journal of Clinical Psychology - Wiley Online Library. 1997;36(4):467–88.

2. Sapolsky RM. The Influence of Social Hierarchy on Primate Health. Science. 2005 Apr 29;308(5722):648–52.

3. Broom M. A Unified Model of Dominance Hierarchy Formation and Maintenance. J Theor Biol. 2002 Nov 7;219(1):63–72.

4. van der Kooij MA, Sandi C. The genetics of social hierarchies. Curr Opin Behav Sci. 2015 Apr 1;2:52–7.

5. Blanchard RJ, Flannelly KJ, Blanchard DC. Life-span studies of dominance and aggression in established colonies of laboratory rats. Physiol Behav. 1988 Jan;43(1):1–7.

6. Ghosal S, Sandi C, van der Kooij MA. Neuropharmacology of the mesolimbic system and associated circuits on social hierarchies. Neuropharmacology [Internet]. 2019 Jan 17 [cited 2019 Apr 8]; Available from: http://www.sciencedirect.com/science/article/pii/S002839081930019X

7. Beiderbeck DI, Reber SO, Havasi A, Bredewold R, Veenema AH, Neumann ID. High and abnormal forms of aggression in rats with extremes in trait anxiety – Involvement of the dopamine system in the nucleus accumbens. Psychoneuroendocrinology. 2012 Dec;37(12):1969–80.

8. Hollis F, van der Kooij MA, Zanoletti O, Lozano L, Cantó C, Sandi C. Mitochondrial function in the brain links anxiety with social subordination. Proc Natl Acad Sci U S A. 2015 Dec 15;112(50):15486–91.

9. van der Kooij MA, Hollis F, Lozano L, Zalachoras I, Abad S, Zanoletti O, et al. Diazepam actions in the VTA enhance social dominance and mitochondrial function in the nucleus accumbens by activation of dopamine D1 receptors. Mol Psychiatry. 2018 Mar;23(3):569–78.

10. Cordero MI, Sandi C. Stress amplifies memory for social hierarchy. Front Neurosci. 2007;1:1.

11. Barik J, Marti F, Morel C, Fernandez SP, Lanteri C, Godeheu G, et al. Chronic Stress Triggers Social Aversion via Glucocorticoid Receptor in Dopaminoceptive Neurons. Science. 2013 Jan 18;339(6117):332–5.

12. Bronson FH, Eleftheriou BE. Adrenal Response to Fighting in Mice: Separation of Physical and Psychological Causes. Science. 1965 Feb 5;147(3658):627–8.

13. Schuurman T. Hormonal Correlates of Agonistic Behavior in Adult Male Rats. In: McConnell PS, Boer GJ, Romijn HJ, Van De Poll NE, Corner MA, editors. Progress in Brain Research [Internet]. Elsevier; 1980 [cited 2019 Apr 8]. p. 415–20. Available from: http://www.sciencedirect.com/science/article/pii/S0079612308600795

14. Timmer M, Sandi C. A role for glucocorticoids in the long-term establishment of a social hierarchy. Psychoneuroendocrinology. 2010 Nov;35(10):1543–52.

15. Weger M, Sevelinges Y, Grosse J, de Suduiraut IG, Zanoletti O, Sandi C. Increased brain glucocorticoid actions following social defeat in rats facilitates the long-term establishment of social subordination. Physiol Behav. 2018 15;186:31–6.

16. Ambroggi F, Turiault M, Milet A, Deroche-Gamonet V, Parnaudeau S, Balado E, et al. Stress and addiction: glucocorticoid receptor in dopaminoceptive neurons facilitates cocaine seeking. Nat Neurosci. 2009 Mar;12(3):247–9.

17. Lemberger T, Parlato R, Dassesse D, Westphal M, Casanova E, Turiault M, et al. Expression of Cre recombinase in dopaminoceptive neurons. BMC Neurosci. 2007 Jan 3;8:4.

18. Pellow S, File SE. Anxiolytic and anxiogenic drug effects on exploratory activity in an elevated plus-maze: A novel test of anxiety in the rat. Pharmacol Biochem Behav. 1986 Mar;24(3):525–9.

19. Kohl C, Riccio O, Grosse J, Zanoletti O, Fournier C, Schmidt M V., et al. Hippocampal Neuroligin-2 Overexpression Leads to Reduced Aggression and Inhibited Novelty Reactivity in Rats. PLoS ONE. 2013;8(2).

20. Wang F, Zhu J, Zhu H, Zhang Q, Lin Z, Hu H. Bidirectional Control of Social Hierarchy by Synaptic Efficacy in Medial Prefrontal Cortex. Science. 2011 Nov 4;334(6056):693–7.

21. Zhou T, Zhu H, Fan Z, Wang F, Chen Y, Liang H, et al. History of winning remodels thalamo-PFC circuit to reinforce social dominance. Science. 2017 Jul 14;357(6347):162–8.

22. Zhou T, Sandi C, Hu H. Advances in understanding neural mechanisms of social dominance. Curr Opin Neurobiol. 2018 Apr 1;49:99–107.

23. Anstrom KK, Miczek KA, Budygin EA. Increased phasic dopamine signaling in the mesolimbic pathway during social defeat in rats. Neuroscience. 2009 Jun 16;161(1):3–12.

24. Fantin G, Bottecchia D. Effect of nucleus accumbens destruction in rat. Experientia. 1984 Jun 15;40(6):573–5.

25. Puciklowski O, Trzaskowska E, Kostowski W, Wośko W. Inhibition of affective aggression and dominance in rats after thyrotropin-releasing hormone (TRH) microinjection into the nucleus accumbens. Peptides. 1988 May 1;9(3):539–43.

26. Bouc RL, Pessiglione M. Imaging Social Motivation: Distinct Brain Mechanisms Drive Effort Production during Collaboration versus Competition. J Neurosci. 2013 Oct 2;33(40):15894–902.

27. Ly M, Haynes MR, Barter JW, Weinberger DR, Zink CF. Subjective Socioeconomic Status Predicts Human Ventral Striatal Responses to Social Status Information. Curr Biol. 2011 May 10;21(9):794–7.

28. Zink CF, Tong Y, Chen Q, Bassett DS, Stein JL, Meyer-Lindenberg A. Know Your Place: Neural Processing of Social Hierarchy in Humans. Neuron. 2008 Apr 24;58(2):273–83.

29. Salamone JD, Correa M. The Mysterious Motivational Functions of Mesolimbic Dopamine. Neuron. 2012 Nov 8;76(3):470–85.

30. Salamone JD, Pardo M, Yohn SE, López-Cruz L, SanMiguel N, Correa M. Mesolimbic Dopamine and the Regulation of Motivated Behavior. In: Simpson EH, Balsam PD, editors. Behavioral Neuroscience of Motivation [Internet]. Springer International Publishing; 2015 [cited 2017 May 30]. p. 231–57. (Current Topics in Behavioral Neurosciences). Available from: http://link.springer.com/chapter/10.1007/7854_2015_383

31. Davis JF, Krause EG, Melhorn SJ, Sakai RR, Benoit SC. Dominant rats are natural risk takers and display increased motivation for food reward. Neuroscience. 2009 Aug 4;162(1):23–30.

32. Kunkel T, Wang H. Socially dominant mice in C57BL6 background show increased social motivation. Behav Brain Res. 2018 Jan 15;336:173–6.

33. Robbins TW, Everitt BJ. A role for mesencephalic dopamine in activation: commentary on Berridge (2006). Psychopharmacology (Berl). 2007 Apr 1;191(3):433–7.

34. Barrot M, Marinelli M, Abrous DN, Rougé-Pont F, Le Moal M, Piazza PV. The dopaminergic hyper-responsiveness of the shell of the nucleus accumbens is hormone-dependent. Eur J Neurosci. 2000 Mar 1;12(3):973–9.

35. Piazza PV, Rougé-Pont F, Deroche V, Maccari S, Simon H, Le Moal M. Glucocorticoids have state-dependent stimulant effects on the mesencephalic dopaminergic transmission. Proc Natl Acad Sci U S A. 1996 Aug 6;93(16):8716–20.

36. Piazza PV, Barrot M, Rougé-Pont F, Marinelli M, Maccari S, Abrous DN, et al. Suppression of glucocorticoid secretion and antipsychotic drugs have similar effects on the mesolimbic dopaminergic transmission. Proc Natl Acad Sci. 1996 Dec 24;93(26):15445–50.

37. Parnaudeau S, Dongelmans M, Turiault M, Ambroggi F, Delbes A-S, Cansell C, et al. Glucocorticoid receptor gene inactivation in dopamine-innervated areas selectively decreases behavioral responses to amphetamine. Front Behav Neurosci [Internet]. 2014 Feb 12 [cited 2015 Oct 2];8. Available from: http://www.ncbi.nlm.nih.gov/pmc/articles/PMC3921555/

38. Larrieu T, Cherix A, Duque A, Rodrigues J, Lei H, Gruetter R, et al. Hierarchical Status Predicts Behavioral Vulnerability and Nucleus Accumbens Metabolic Profile Following Chronic Social Defeat Stress. Curr Biol. 2017 Jul 24;27(14):2202–2210.e4.

39. Gilbert P, McEwan K, Bellew R, Mills A, Gale C. The dark side of competition: How competitive behaviour and striving to avoid inferiority are linked to depression, anxiety, stress and self-harm. Psychol Psychother Theory Res Pract. 2009 Jun 1;82(2):123–36.

40. Goette L, Bendahan S, Thoresen J, Hollis F, Sandi C. Stress pulls us apart: Anxiety leads to differences in competitive confidence under stress. Psychoneuroendocrinology. 2015 Apr 1;54:115–23.

