## Supplementary information for "The glucocorticoid receptor in the nucleus accumbens plays a crucial role in social rank attainment in rodents"

### Supplementary Materials and Methods

##### Elevated plus maze test

The apparatus consisted of two opposing open arms perpendicular to two enclosed arms (50 x 10 x 50 cm) that extend from a central platform (10 x 10 cm), elevated 65 cm above the floor. Light levels were maintained at 14-16 lux on the open arms and 5-7 lux in the closed arms. At the start of the test, the animal was placed on the central platform facing a closed arm and allowed to explore the maze for five minutes. In between animals, the apparatus was cleaned with 5% ethanol solution and dried thoroughly between each animal. Behavior was monitored using a ceiling-mounted video camera and analyzed with a computerized tracking system (Ethovision 9, Noldus IT, Netherlands). The time spent in the open and closed arms and distance moved were automatically recorded.

##### Open field and novel object reactivity tests

A circular open arena was utilized (1 m diameter, 40 cm high). Each rat was placed near the wall and it was allowed to explore the apparatus freely for 10 minutes. The floor of the open field was virtually divided in three virtual parts: center zone in the middle of the arena with a diameter of 25 cm, an intermediate zone with a diameter of 75 cm and the remaining wall zone along the walls of the arena. At the end of the 10-minute period, an object (yellow plastic bottle) was introduced into the center of the arena for the novel object reactivity test. Each animal was allowed to explore the arena and object for 5 minutes. The time spent in each zone, the total distance moved and the time spent sniffing the object were recorded. The apparatus was cleaned with 5% ethanol solution and dried thoroughly between each animal.

**Social hierarchy test**

The test was performed as described previously (8). Rats were pair-wise matched for weight but animals in each pair were of opposite anxiety profile. Animals were marked on their body for identification and placed in pairs in a clean (neutral) cage without food or water for 20 minutes. During the social hierarchy test both rats displayed spontaneously offensive behavior, but this balance typically shifted in a favor of one animal towards the end of the test. The following parameters were quantified in terms of duration: offensive upright, lateral threat and keeping down. The cumulative duration of these behaviors was summed to provide a measure of total offensive behavior. Pairs in which offensive behavior was virtually absent during social encounter (no rat displaying > 10s of total offensive behavior) were excluded from the analysis, as the relative social dominance in these pairs cannot be reliably measured.

**Social confrontation tube test**

The protocol was adapted from (20). For the rat experiments, the tube test was performed in dyads that had been living together for 2 weeks. Each rat was individually trained to move forward out of a clear Plexiglas tube (diameter, 7.5 cm; length 100 cm) over five consecutive days, with two trials per day. Each rat of the dyad learned to move forward only from one end. The size of the diameter is just sufficient to permit an adult rat to move through the tube without reversing its direction. If the rat retreated or stopped moving for a certain amount of time, it was gently pushed by touching its tail with a plastic stick. The tube was cleaned and dried between each trial with 5% ethanol to remove odor, urine or feces. After five days of training, dominance was evaluated for five consecutive days. The two rats of each dyad were handled and guided simultaneously at the opposite ends of the tube until they entered the tube and reached the middle part. The time spent in the tube was recorded until one of the two rats forced its cage mate to go backwards and exit the tube. The rat that retreated from the tube was designated as the ‘loser’ of that trial. The rat that won four trials or more was considered dominant and the loser was the subordinate rat.

The tube test protocol on mice was similar to that performed in rats, except that the dyads had been living together for one week before being tested. The apparatus was smaller (diameter, 3cm; length 30cm) and mice were trained over seven consecutive days, with three trials per day. The dominance was then evaluated for three consecutive days, with three trials per day. The mouse that won seven trials or more was the dominant and the loser was the subordinate mouse.

**Water competition test**

To corroborate the social hierarchy indexed in the tube test in the home cage, we performed water competition test. Rats were water-deprived for 6 hours, before the beginning of the test. After marking the fur of both rats in a dyad to help identify the animals, a single bottle of water was presented and the time spent drinking during the following 10 minutes was recorded. The animal that spent the greater percentage of time drinking the water was considered the dominant of the dyad.

**Warm spot test**

The mice were placed in the experimental room one hour before the experiment for habituation. They were marked with different colors on the tail in order to discriminate between the animals on the video. Two empty cages (28 cm x 20 cm, without litter), A and B, were placed on ice until their floor temperature reached 0°C. A heating mat (Thermo Mat, 3W, Lucky Reptile, about 38°C) was inserted through a notch in one corner of the cage A (size 4 cm x 3 cm, so that only one mouse at a time could access the mat). The two mice of the dyad were put in cage B (on ice, without heating mat) for 20 min and were then transferred to cage A for another 20 min. The session was video recorded and the time of mat occupancy was measured for each mouse. The mouse who spent the greater percentage of time on the mat was considered the dominant of the dyad.

**Viral downregulation of GR**

The vector incorporated the following regulatory elements: rAVE™ construct containing EGFP, the hybrid chicken B-actin/CMV enhancer (CAG) promoter region, a cis-acting woodchuck post-transcriptional regulatory element (WPRE) and a bovine growth hormone polyadenylation sequence (BCG-polyA). A perfect match (pm) short hairpin RNA (shRNA) construct against GR, driven by U6 promoter was also incorporated. For the scrambled vector, the same backbone without the cDNA was used. Initially, the animals were anaesthetized by inhalation of isoflurane (5% oxygen-isoflurane mixture) and installed in a stereotaxic frame to avoid any head movements during the surgical procedure. To target the nucleus accumbens, two injections sites were used with the following coordinates (42): 1.3 and 2.5 mm posterior to bregma, 1.0 and 1.5 mm from midline, 7.0 ventral from skull. A volume of 0.8 µl of either GR-KD or scrambled vector (titres: $1.3x10^{12}$ genomic particles/ml) was bilaterally injected in the nucleus accumbens with a constant flow rate of 0.1 µl/min. The injectors were left in site for 5 minutes after the end of the actual injection. After removing the injectors, animals were treated with paracetamol (500 mg/700 ml H2O, Dafalgan, Bristol-Myerts

Squibb, Agen, France) via the drinking water for seven days after the surgery. The animals were allowed to recover for 6 weeks from surgery so that the maximum viral expression could be reached before the behavioral testing was started.

**Gene expression analysis, immunofluorescence and immunohistochemistry**

Rats were decapitated under basal conditions. After decapitation, brains were snap frozen in isopentane at -45°C and stored at -80°C. They were then sectioned using a cryostat and 200 µm thick slices were mounted on slides in order to punch and remove the nucleus accumbens (NAc), with a tissue puncher of 1 mm. The tissue punches were collected in RNAse-free tubes. Total RNA from the Nac was isolated using the RNAqueous Micro kit (Ambion, Life Technologies, USA), and complementary DNA was synthesized using the qScript cDNA Mastermix kit (Quanta Biosciences, USA) according to supplier's recommendations. For real-time quantitative polymerase chain reaction (qPCR), reactions were performed in triplicate using SYBR Green PCR Master Mix (Applied Biosystems, Life Technologies, USA) in an ABI Prism 7900 Sequence Detection system (Applied Biosystems, Singapore). Two genes were used as internal controls: actin gamma 1 (ActG1) and eukaryotic elongation factor 1 (EEF1). We analyzed the expression of the glucocorticoid receptor (GR) and the mineralocorticoid receptor (MR). Primers for the genes of interest were designed using the Assay Design Center software from Roche Applied Science (Table S1). Gene expression was analyzed with using the Pfaffl method (PFaffl MW, A new mathematical model for relative quantification in real-time RT-PCR), and normalized against the geometric mean of the expression of the internal controls.

| Gene | Full name | Forward primer (5'-3') | Reverse primer (5'-3') | RefSeq (NCBI) |
|---|---|---|---|---|
| **EEF1** | Eukaryotic translation elongation factor 1 | tgtggtggaatcgacaaaag | cccaggcatacttgaaggag | NM_175838.1 |
| **ActG1** | Actin gamma 1 | tagttcatgtggctcggtca | gctggggactgactgacttt | NM_001127449.1 |
| **GR** | Glucocorticoid receptor | aaagcttctggactccatgc | tcaatactcatggtcttatccaaaaa | NM_012576.2 |
| **MR** | Mineralocorticoid receptor | caagctggcatgaacttagga | tcctcgtggaggcctttt | NM_013131.1 |

**Table S1. Primer sequences for real-time qPCR.**

In the GR knockdown experiment, one brain hemisphere was used for gene expression analysis. The second hemisphere was fixed in PFA 4% for 48 hours before being cryoprotected in 30% sucrose solution and frozen at -80°C. Subseries of coronal sections (30 µm thick), including the nucleus accumbens, were cut on a cryostat and processed for immunofluorescence or immunohistochemistry. In order to validate the viral delivery, free-floating sections were double labeled for GR, DAPI and the green-fluorescent protein (GFP) expressed by the virus. The floating sections were rinsed briefly with PBS containing 0.5% Triton X-100 (Sigma-Aldrich), blocked for 1.5

hours in PBS containing 0.5% Triton X-100 and 5% normal donkey serum (Jackson ImmunoResearch), and then incubated overnight at 4°C with rabbit anti-GR (Santa Cruz, M-20, 1:1000). The sections were washed in PBS and incubated for 2 hours at room temperature with the secondary antibody: donkey-anti-rabbit IgG Alexa 568 conjugate (Life Technologies, A10042, 1:800). After washing in PBS, the sections were incubated 10 minutes in DAPI (Sigma, 1:10000), rinsed and mounted with Fluoromount-G (Southern Biotech). Images were captured with an Olympus Slide Scanner VS120-L100 using a 20x objective.

Efficiency of virally-mediated GR knockdown was tested by DAB immunohistochemistry. After incubation in 0.3% $H_2O_2$/PBS to block endogenous peroxidases, the free-floating sections were blocked in 10% donkey serum/PBS containing 0.2% Triton X-100 (PBS-T) and incubated overnight with the primary antibody against GR (Santa Cruz, M-20, 1:500) at 4°C. Sections were rinsed in PBS-T and then incubated with the secondary antibody (biotinylated anti-rabbit IgG, Vector Laboratories USA, 1:1000) for 2 hours. Afterwards, sections were treated using an ABC kit (Vectastain ABC Kit, Vector Laboratories, USA) followed by further washes in PBS until color development using 3,39-diaminobenzidine (DAB substrate kit for peroxidase, Vector Laboratories). Images were taken with the Olympus Slide Scanner VS120-L100 using a 40x objective. Two animals injected with the scrambled construct were excluded from the analysis due to a wrong localization of the virus as indicated by immunofluorescence.

#### Supplementary Results

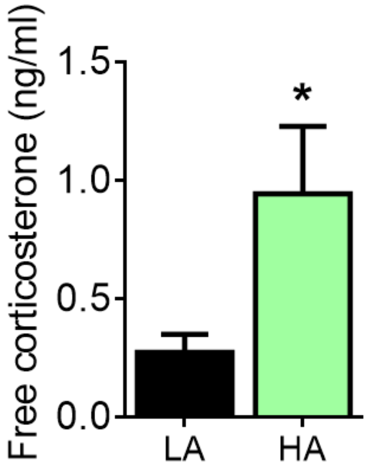


**Figure S1. High anxious rats exhibit higher levels of basal corticosterone than low anxious counterparts.** High anxious (HA; equivalent to submissive-prone, subP) rats had increased corticosterone at baseline compared to low anxious (LA; equivalent to dominant-prone, domP) conspecifics ($U$ = 9.50, $p < 0.05$). *N:* LA = 7, HA = 8. *$p < 0.05$, vs SCR. Results are expressed as mean ± SEM.

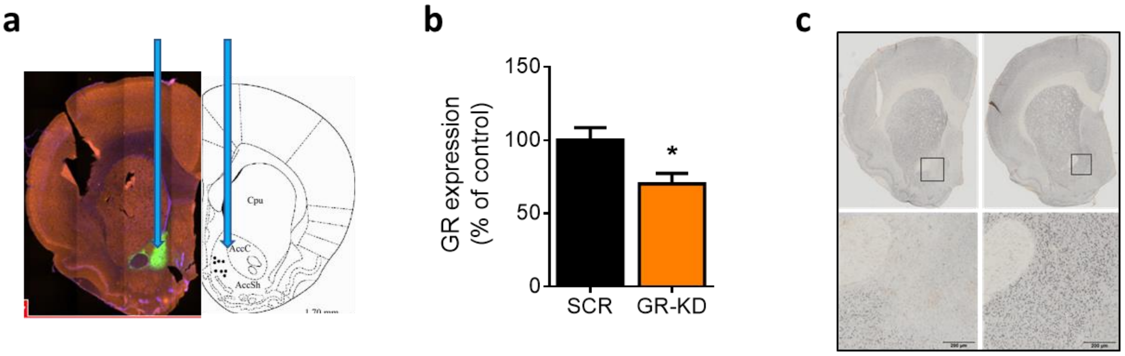


**Figure S2. Verification of viral GR downregulation in the nucleus accumbens.** Targeted area in the NAc and GFP localization with 4x magnification of the NAc are shown in **(a)**. GR mRNA expression in the NAc was decreased in rats injected with the AAV-U6-shGR virus (GR-KD) compared to scrambled (SCR)-infused rats ($U$ = 4.00, $p$ < 0.05) **(b)**. DAB immunohistochemistry (40x magnification) further confirmed the downregulation of GR in the nucleus accumbens in GR-KD animals (left panel) compared with SCR rats (right panel) **(c)**. *N:* SCR = 6; GR-KD = 6. *$p$ < 0.05, vs SCR. Results are expressed as mean ± SEM.

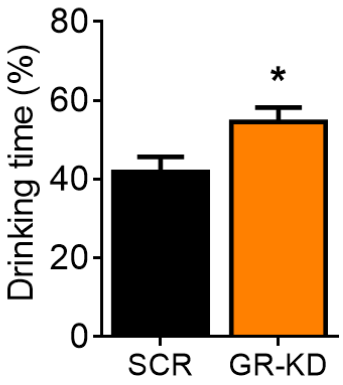


**Figure S3. Water competition in GR-KD rats.** The water competition test confirmed the dominance determined by the social confrontation tube test, showing that GR-KD rats spent more time drinking than SCR conspecifics ($t_{22}$ = 2.40, $p < 0.05$). *N:* SCR = 12, GR-KD = 12. *$p < 0.05$, vs SCR. Results are expressed as mean ± SEM.

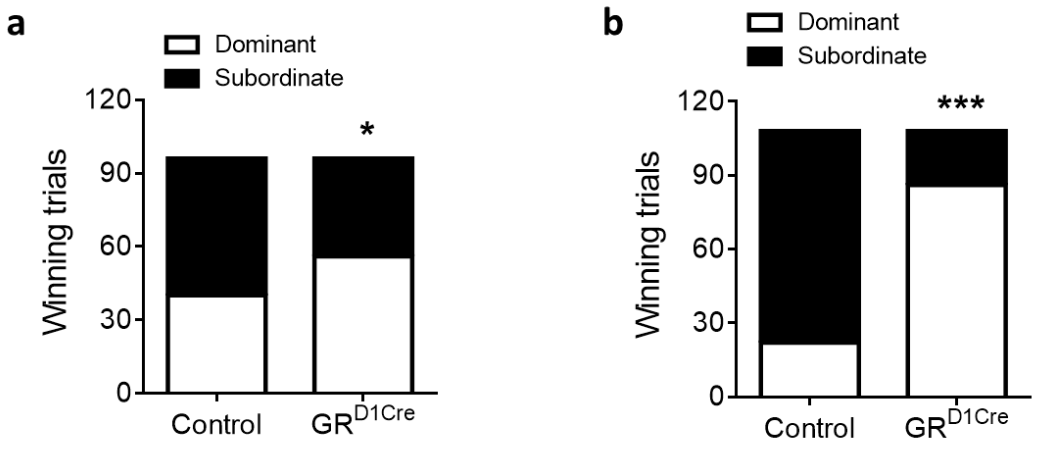


**Figure S4. Tube test analysis from two different experiments.** $GR^{D1Cre}$ mice showed higher number of winning trials than control animals (Fisher's exact test, $p < 0.05$) **(a)**, which has been replicated a second time in an independent experiment (Fisher's exact test, $p < 0.001$) **(b).** N: First experiment: control = 11, $GR^{D1Cre}$ = 11; second experiment: control = 12, $GR^{D1Cre}$ = 12. *$p < 0.05$, ***$p < 0.001$, vs control. Results are expressed as mean ± SEM.

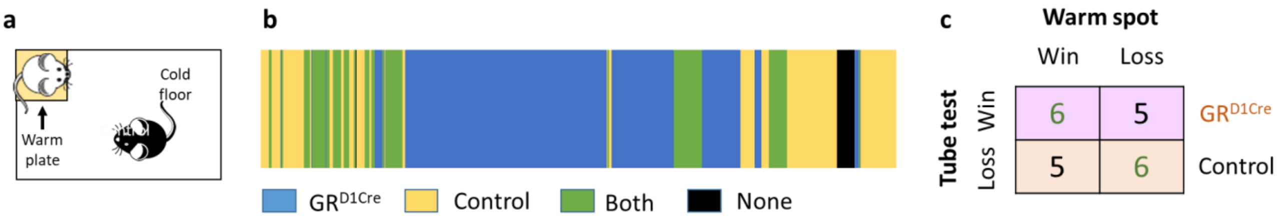


**Figure S5. Warm spot test in mice lacking GR in NAc dopaminoceptive neurons**. Schematic of the warm spot test in which a warm plate is placed in one corner of a cold cage. The time spent by each mouse on the warm spot is used as a measurement of dominance **(a)**. Example of occupancy of the warm spot within a dyad **(b)**. Correlation between the tube test and the warm spot test. Out of 11 dyads, six $^{GRD1Cre}$ mice won their contests in both the tube test and the warm spot test **(c)**. N: Control = 11, $GR^{D1Cre}$ = 11. Chi square test: $\chi^2 = 5.12 \geq \chi^2$ 1.5% = 3.84. *$p < 0.05$, vs control.

#### Supplementary Reference

41. Paxinos, G., and Watson, C. The rat brain in stereotaxic coordinated: hard cover edition. Academic Press. 2006
